# An integrated mathematical model of the neuromuscular activity of a motor unit

**DOI:** 10.1101/2023.12.06.570328

**Authors:** Zdravka D. Ivanova, Tihomir B. Ivanov, Rositsa T. Raikova

## Abstract

In the present work, we propose a new integrated mathematical model for the neuromuscular activation of a motor unit, describing the four consecutive processes, leading to muscle contraction—neural impulse propagation, acetylcholine transport in the neuromuscular junction, calcium release in the muscle cell, and force generation. We connect in an appropriate way models of the respective processes, known from the literature, and validate the resulting model by showing that it can reproduce with high accuracy experimental data for two motor unit twitches on a rat medial gastrocnemius muscle and can numerically restore the sequence of events that result in force generation. Sensitivity analysis for some of the model parameters is further performed to study their effect on the model solutions and to show that they can be related to known malfunctions or treatments of the neuromuscular system.

## 1. Introduction

The excitability of *α*-motoneurons and muscle cells and their corresponding behaviour is essential for the understanding of muscle contraction and neuromuscular diseases (NMDs).

On one hand, mathematical models, capable of accurately simulating muscle contraction, are very useful not only for the better understanding of the underlying processes, but also for numerous bioand biomedical engineering applications. However, the precise mathematical description even of a single motor unit (MU) twitch, i.e., a single contraction of the MU—the basic functional unit in the skeletal muscle, has shown to be a very challenging task. There exist phenomenological models [1, 2], which are capable of reproducing the shape of a MU twitch (as shown from experimental data), but to the best of the authors’ knowledge **there are no descriptive models, which have been fitted with a high level of accuracy to experimental data for a MU twitch** [2].

On the other hand, NMDs affect the normal functioning of the muscles in human body and are connected with a broad group of disorders in the MUs. The different NMDs affect different sites of the MU—the nerve, neuromuscular junction (NMJ), muscle cells or combination of those. A more complete description of a MU can be found, e.g., in [3, 4, 5]. Therefore, **in order to study NMDs, one needs to be able to study how the characteristics of each part of the MU affect its behaviour**. In Ivanova and Ivanov (2022) [6], some results were obtained for the dependence of the generated muscle force on various firing patterns and properties of the muscle cell. In the work [6], however, a biologically plausible description of the NMJ was not included. Nevertheless, understanding the effect of its properties and their relation to the process is extremely important for various NMDs.

Therefore, in the present paper, we aim:

- to construct a mathematical model of neuromuscular activation, described in terms of differential equations, which can successfully reproduce MU twitch data;
- to perform sensitivity analysis, i.e., to show how some of the main parameters in the model, related to the properties of the NMJ, affect the shape of the MU twitch.

Let us begin with a brief description of the main processes, involved in the MU activation. *α*-motoneurons innervate the skeletal muscles to contract (see Fig. 1). When electrically stimulated, the function of the skeletal MUs is to produce force. Muscle force is the sum of forces of all activated MUs. The processes behind the activation of a MU can be summarized as follows [7]:

**Figure 1:**
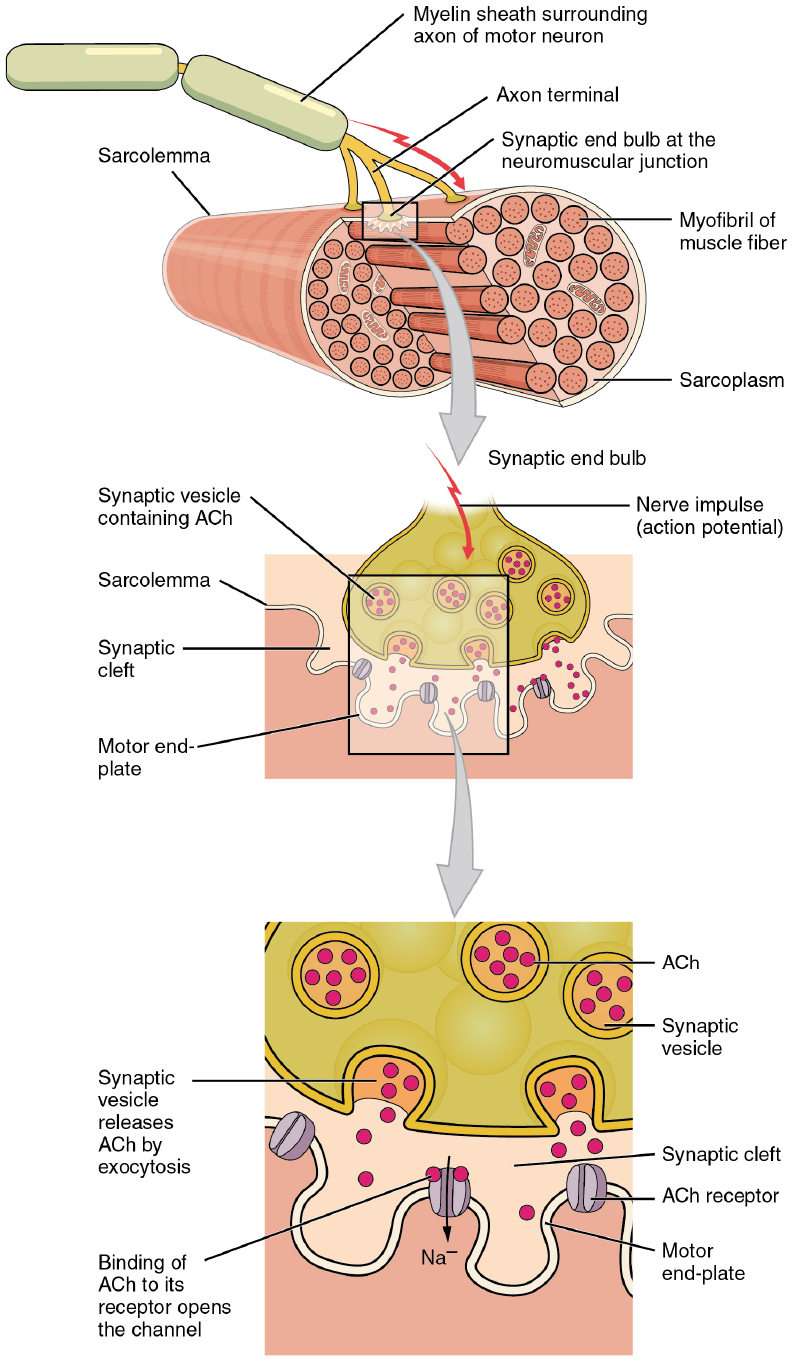
Coupling between the nerve and muscle cells (the image is reproduced from [8], available under CC BY 4.0 license).

### 1. Action potential and release of neurotransmitter

An impulse, called action potential, travels through the axon of the motor neuron to the axon terminal, resulting in a release of acetylcholine (ACh) in the neuromuscular junction. ACh then binds to the receptors on the motor end-plate.

### 2. Calcium release in the muscle fiber

The receptors at the end-plate of muscle fiber open and sodium ions enter into the muscle cell, causing potassium ions to exit it. The input flux of the sodium ions changes the membrane potential, causing depolarization or the so-called end-plate potential. This depolarization can cause a depolarization of the muscle fiber once the membrane potential reaches a threshold value. Further, an action potential propagates into the interior of the fiber and leads to calcium release from the sarcoplasmic reticulum (SR).

### 3. Calcium binds to contractile filaments

The released calcium can bind to the actin filaments in the muscle fiber. The binding of calcium allows the contractile filaments (CFs)—actin and myosin filaments, to bind to each other, forming a structure called a cross-bridge, and further leads to muscle cell contraction. After the cross-bridge formation, calcium is pumped back into the SR, lowering the calcium level in the sarcoplasm and leading to relaxation.

Thus, the process of MU contraction can be summarized as a result of four different consecutive processes—propagation of nerve impulses, neurotransmitter release and transport in the NMJ, the resulting biochemical reactions in the muscle cells, and the generated contraction. To that end, basically, we shall consider a more complete implementation of the general scheme, shown in Fig. 2, which we introduced in [6].

**Figure 2:**
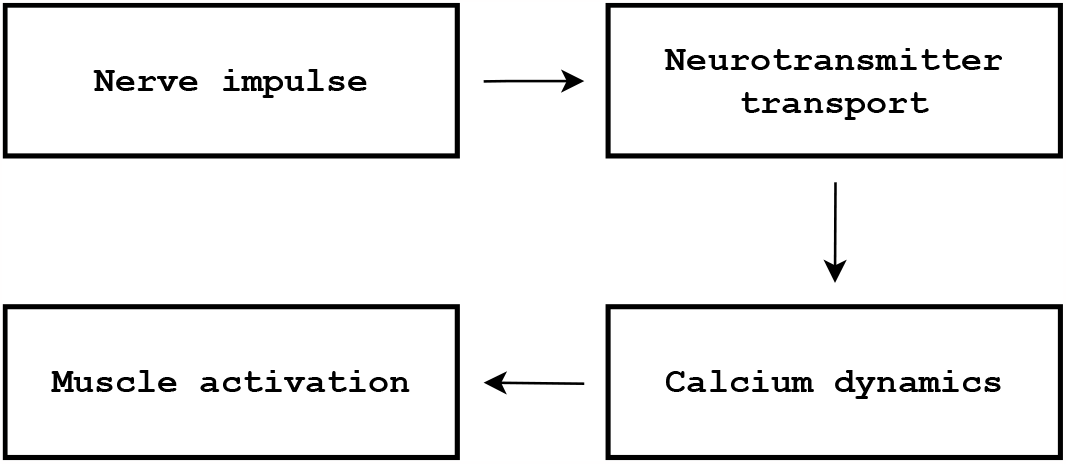
General scheme for the modelling of the process of muscle activation.

To obtain a model of neuromuscular activation, capable of showing how variations in its properties affect the MU behaviour, one should apply a detailed integrated approach, in particular modelling the steps in the latter scheme. Unfortunately, in the scientific literature such kind of models are extremely scarce [6, 9, 10, 11]. On the other hand, each individual process (or sometimes a couple of processes) has been the subject of various works. Let us mention some of them.

- Nerve impulse modelling—there exist plenty of mathematical models, having various degrees of complexity and biological plausibility. On the two ends of the spectrum are the classical models of Hodgkin–Huxley [12] and McCulloch–Pitts [13], which are, respectively, a detailed biological description and a very simplistic description of the process. Using various simplifications of the Hodgkin–Huxley model, taking into account its high computational complexity, one may consider models, such as the Fitzhugh–Nagumo model [14, 15], the Hindmarsh–Rose model [16], the Morris–Lecar model [17], the Markin–Chizmadzhev model [18], etc. The Izhikevich model [19] offers complex dynamics that allows for simulating many different nerve firing patterns. It is relatively simple and is, thus, of particular importance for our purposes. A more detailed review of various mathematical models of neural activity along with some of their properties can be found, e.g., in [20, 21, 22] and the references therein. Those models can either be employed as systems of ordinary differential equations (ODEs) to simulate the temporal behaviour of the neural activity or can be combined with the cable equation (or its discrete counterpart for myelinated nerves) for a more detailed spatio-temporal description of the axonal transport (see, e.g., [22, 23] and the references therein);
- Neurotransmitter transport in the neuromuscular junction—a 2D reaction-diffusion model was proposed by Naka et al. in [24], which was further extended by Khaliq et al. in [25]. Finite element simulations of the ACh diffusion in the synaptic cleft, based on a continuum mechanics description, were performed, e.g., in [26];
- Calcium dynamics inside the muscle cell—there exist reaction-diffusion models, such as the model described in [27], as well as ODE models, only describing the temporal dynamics of the main chemical reactions such as the one, proposed by Williams et al. in [28, 29];
- Muscle force—we can divide those models to phenomenological and physically-based (with various degree of detailedness). Phenomenological models include the ones by Fuglevand [30] and Raikova et al. in [1, 31]. A more complete review on the subject can be found in [2]. Classical physically-based models include the models of Hill [32] and Huxley [33]. Some information about such kind of models can be found in, e.g., [34, 35]. A continuum mechanics based approach can be found, e.g., in [10, 36, 37] and the references therein.

Therefore, we aim to base our work on known results for the various processes as well as on the ideas, which we proposed in [6], and make one step further towards obtaining an integrated mathematical model of the complete process. For this work, we shall couple the following models for each step, for reasons that we describe below:

- Izhikevich model for the action potential [19]—this model is chosen because it combines simplicity with very complex dynamics, allowing for the simulation of many different firing patterns;
- A model of the ACh transport in the NMJ, proposed by Naka et al. [24];
- A model proposed by Williams [29] of the calcium dynamics inside the muscle cell—again, as a first step in this direction, we make a choice in favour of simplicity; furthermore, the model has been validated in terms of real experimental data;
- Hill-type model for the muscle contraction [32]—these are classical (yet simple) models, which have been employed extensively in the scientific literature; furthermore, a Hill-type model has been coupled with the Williams model in [29]; in the present work, we revisit this coupling by incorporating some ideas, proposed in [27] and [38].

Let us re-emphasize that the model, which we formulate, should serve as a step further in the construction of a hierarchy of mathematical models with various degree of detailedness, for which we laid the foundation in [6], and which would aim to shed light on different aspects of the process. Certainly, one can argue that another choice of models for each step could be more appropriate, but our aim is to start with a model that suits our requirements and, then, go on to make it ever more biologically plausible. Therefore, we are mainly interested here in the general approach and in the types of results we could obtain, using such an approach.

The main novelty, which we introduce with respect to the model, we considered in [6], is the detailed description of the ACh transport in the NMJ. This is a very important improvement of the model, because this was the missing connection between all the stages of the process. Another improvement is the introduction of an activation function, relating the calcium dynamics inside the muscle cell with the resultant MU force. As we show in the present work, the improved model is now capable of fitting experimental data for a MU twitch with a high degree of accuracy. We validate the applicability of the model against real MU twitch data and we further study how the properties of the NMJ affect the solutions of the model. We focus here on this aspect of the complete process, since, as we have noted, describing the NMJ is the main novelty with respect to [6].

The present work is structured as follows. In Section 2, we formulate the mathematical model, by giving the models we use for each stage of the process and explain how we couple them. In Section 3, we validate the model by showing that it can successfully reproduce experimental data for two MU twitches by choosing appropriate values of the model parameters. In Section 4, we perform sensitivity analysis to study how varying the model parameters, corresponding to the properties of the NMJ, affects the characteristic parameters of the twitch. Section 5 is a summary and discussion of the obtained results.

## 2. Mathematical formulation

Let us first formulate here the Izhikevich model of neural impulse [19]:

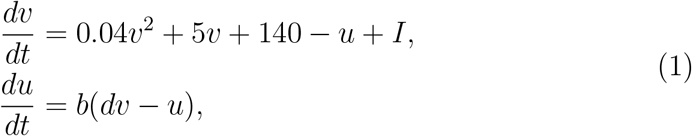

with the auxiliary rule that if *v* = 30*mV* holds, then, *v ← v*_*reset*_, *u ← u* + *u*_*reset*_, i.e., the variables *v* and *u* are reset after each spike (or, equivalently, when *v* gets to 30mV) to values *v*_*reset*_ and *u* + *u*_*reset*_, correspondingly. Here, *v*(*t*) means the transmembrane voltage at time *t* and *u*(*t*) is a recovery variable. The parameters *b, d, u*_*reset*_, *v*_*reset*_ are positive constants and *I* = *I*(*t*) is the input current at time *t*.

The main novelty with respect to the model, which we consider in [6], is the detailed description of the ACh transport in the NMJ (instead of just coupling directly the model of neural activity (1) and the resulting calcium activity as we did in [6]). To model the ACh transport in the synaptic gap, we consider a model proposed by Naka [24] and make further an assumption to consider an ODE system.

When an action potential reaches the end of the axon, a neurotransmitter called acetylcholine is released in the synaptic cleft. ACh then diffuses to the post-synaptic membrane, where it binds to special receptors (R). In the process, the ACh reacts with the enzyme acetylcholinesterase (AChE), which catalyzes the breakdown of ACh into choline and acetate. Further, on the post-synaptic membrane, when doubly bound with ACh, the receptors open, allowing for the impulse to “go into” the muscle cell (see also Fig. 1). The process can be described by the following reaction schemes (see [24], [25], and [39], and the references therein):

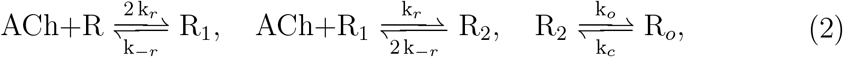

where *R* and *R*_1_ denote the free and single-bound receptors, *R*_2_ denotes the double bound closed ACh receptors, and *R*_*o*_ denotes the open ACh receptors; *k*_*r*_, *k*_*−r*_, *k*_*o*_ and *k*_*c*_ are rate constants for binding, unbinding, opening and closing of the specified receptors, respectively.

The reaction of ACh with AChE can be described by the following reaction scheme:

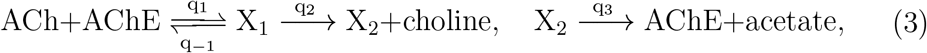

where *X*_1_ and *X*_2_ denote AChE complexed with ACh and acetyl group, respectively; *q*_1_, *q*_*−*1_, *q*_2_ and *q*_3_ are rate constants. Choline and acetate are end products of the reactions and will not be considered further in the models.

To model the process, let us denote the following:

- *a*(*t*)—concentration of ACh at time *t*;
- *r*_1_(*t*), *r*_2_(*t*), *r*_*o*_(*t*)—concentrations of single-bound (*R*_1_), double-bound (*R*_2_), and opened (*R*_*o*_) receptors, respectively, at time *t*;
- *x*_1_(*t*), *x*_2_(*t*)—concentrations of AChE complexes *X*_1_, *X*_2_ at time *t*.

By applying the Law of mass action [40] for the reaction schemes (2), (3), we consider the following model [24] with the assumption that the ACh molecules travel fast in the neuromuscular gap and, thus, we neglect the spatial terms.

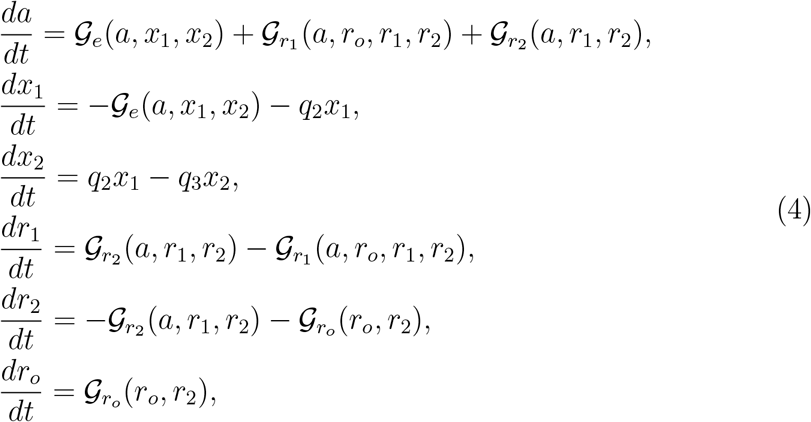

where

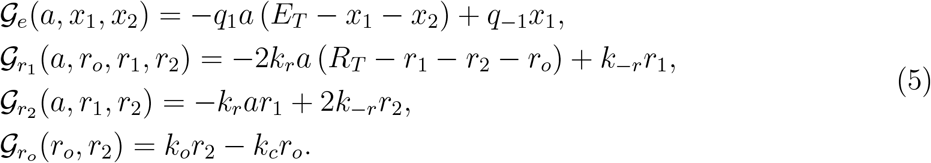

In the latter equations, we have used the following notation: *R*_*T*_ and *E*_*T*_ are the total concentrations of receptors and AChE, correspondingly, and *k*_*r*_, *k*_*−r*_, *k*_*o*_, *k*_*c*_, *q*_1_, *q*_*−*1_, *q*_2_, *q*_3_ are the positive rate constants, given in (2) and (3).

We model the release of ACh and make connection with the nerve impulse signal in the following way. At the times of the local maxima of *v* (i.e., the times of the neural impulses) we assign *a ← a* + *a*_*quant*_, where *a*_*quant*_ is the amount of released ACh (i.e., at those moments of time, we increase the concentration *a* with *a*_*quant*_).

When ACh receptors open, this leads to an input electrical signal in the muscle cell, which in turn causes the release of calcium ions from the SR. We shall assume for simplicity that the stimulus of the muscle cell is proportional to the concentration of open ACh receptors, *r*_*o*_. The released calcium then binds to the CFs, which leads to a muscle contraction. In order to model the calcium dynamics and the resultant MU force, we modify the model, proposed by Williams [29] and used in [9] and [6]. It can be reduced to the following system of 3 ODEs after proper rescaling of the model parameters (for more details, see [41], where we have also studied the properties of the model solutions of the original Williams model in two important limiting cases).

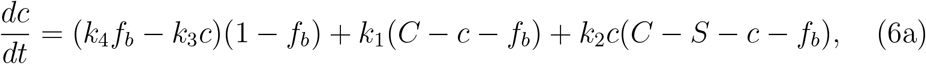

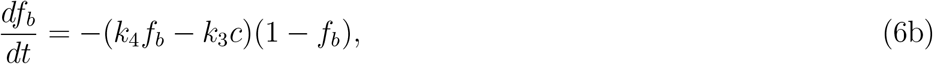

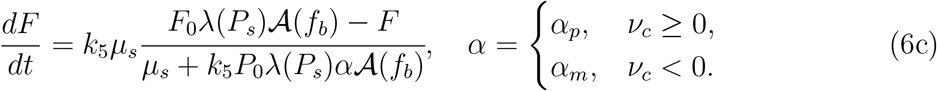

In the latter model, *c*(*t*) is the nondimensionalised concentration of free calcium ions inside the muscle cell at time *t*; *f*_*b*_(*t*) is the nondimensionalised concentration of bound CF sites at time *t*; *F* is the resultant MU force; *k*_1_, *k*_2_, *k*_3_, *k*_4_ are the rates of release of calcium ions from the SR, binding of calcium ions to the SR, binding of calcium ions to the CFs, and rate of release of calcium ions from the CF, correspondingly; *C* and *S* are the total nondimensionalised amounts of calcium and SR sites per unit volume. The concentrations in the original model have been scaled with the total amount of CF sites per unit volume.

For the rates of binding and unbinding from the SR in the mathematical model the following is valid:

- the parameter *k*_1_ is directly proportional with a coefficient *k*_*p*_ *>* 0 to the concentration of open receptors, *r*_*o*_:

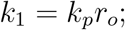

- the parameter *k*_2_ is chosen to be

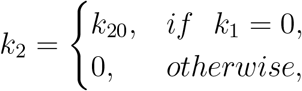

where *k*_20_ is a positive parameter.

Thus, when a stimulus is present *k*_1_ *>* 0, *k*_2_ = 0, and vice versa.

Further, *k*_3_, *k*_4_, *k*_5_ are positive constants; *l*_*s*,0_ and *l*_*c*,0_ are the resting lengths of the spring and contractile elements (in the sense of a Hill-type model [32]), correspondingly; *F*_0_ is the peak force in an isometric tetanic contraction at the optimum length *l*_*c*,0_; *L* is the muscle length; *µ*_*s*_ is the stiffness coefficient;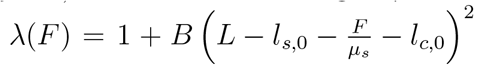. The difference with the original model of Williams and the model, which we consider in [6], lies in the fact that we scale *f*_*b*_ with a sigmoid function to define the activation function *𝒜*(*f*_*b*_) (instead of just using *f*_*b*_ as in the original model) to be

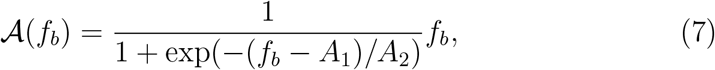

where *A*_1_, *A*_2_ are positive constants.

The reason for doing so lies in the fact that our numerical experiments have shown that such a modification is necessary if one wants to obtain a good fit with respect to experimental data, as we do in the next section. It seems that there must be some minimal concentration of *f*_*b*_ before the MU can start exerting some positive force. The idea, motivating the introduction of such an activation function, however, is not new. Kim et al. [27] state, based on an earlier work [38] that cooperativity of cross-bridge formation has been characterized as a sigmoidal relationship between the sarcoplasmic *Ca*^2+^ concentration and force production under steady-state conditions and further elaborate on this idea in their model. Our comparison to experimental data in the following section shows that the form (7) of *𝒜* (*f*_*b*_) captures the idea sufficiently well for our purposes.

The general coupling between the models, which we presented above, is illustrated schematically in Fig. 3.

**Figure 3:**
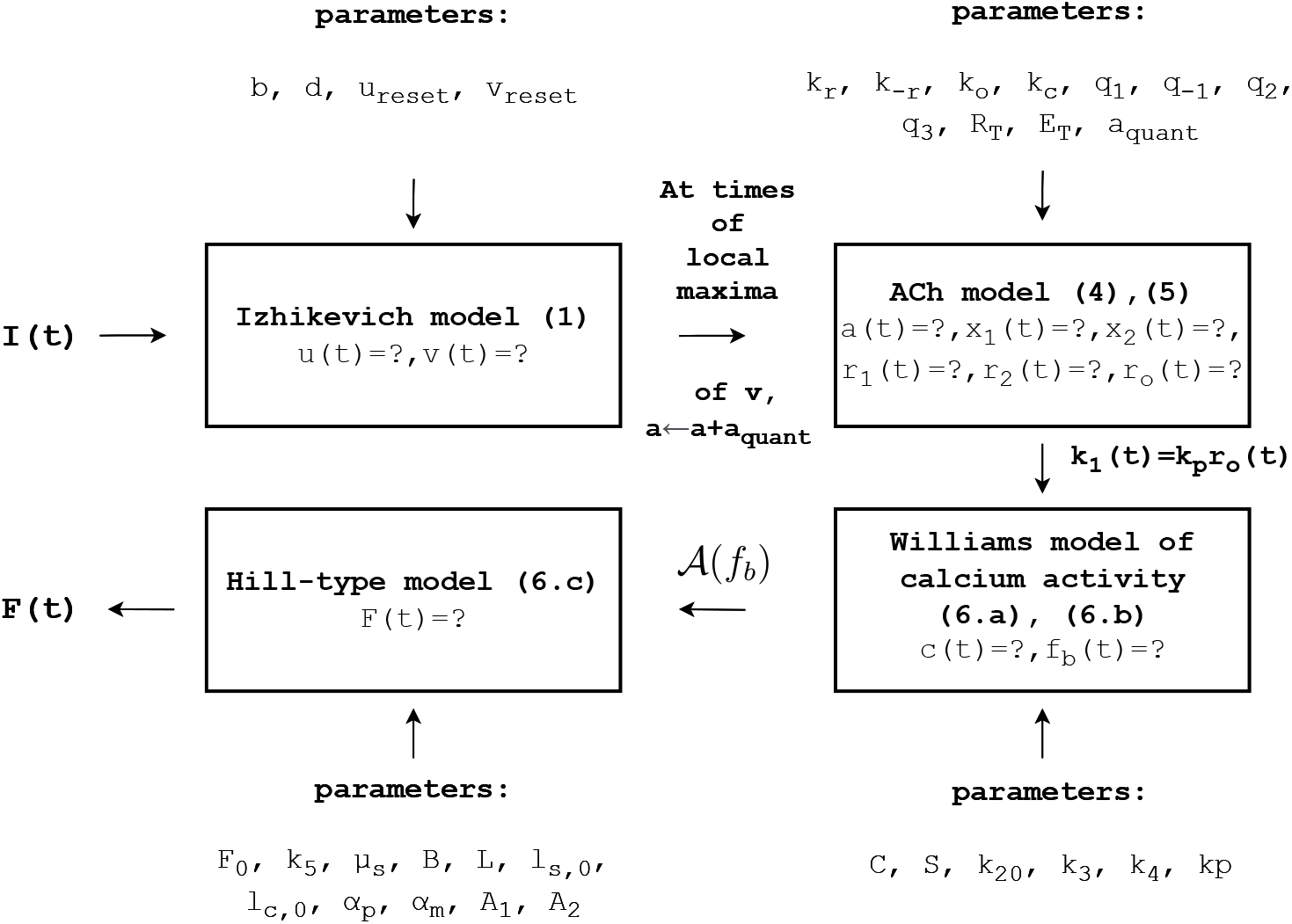
Schematic representation of the computational flow in the integrated mathematical model of neuromuscular activation.

In what follows, we are interested in showing that our model successfully recovers the shape of an experimentally recorded twitch and, furthermore, to study how the different parameters in the model, corresponding to the properties of the NMJ (the main novelty with respect to [6]), affect the twitch (quantified in terms that we define below).

## 3. Validation of the model on a single twitch

In order to validate the model (1), (4), (6), we use data for two different MU twitches on rat medial gastrocnemius, which has been provided to us from the lab of Prof. Jan Celichowski from Poznan University, Poland. The experimental protocol is described in [42]. We are given two sets of data points for the MU twitch curves, having the form

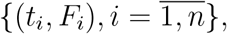

where *F*_*i*_ is measured MU force at time *t*_*i*_, respectively—those are the dots in Fig. 4. We define the following objective function

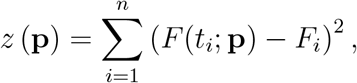

where

**Figure 4:**
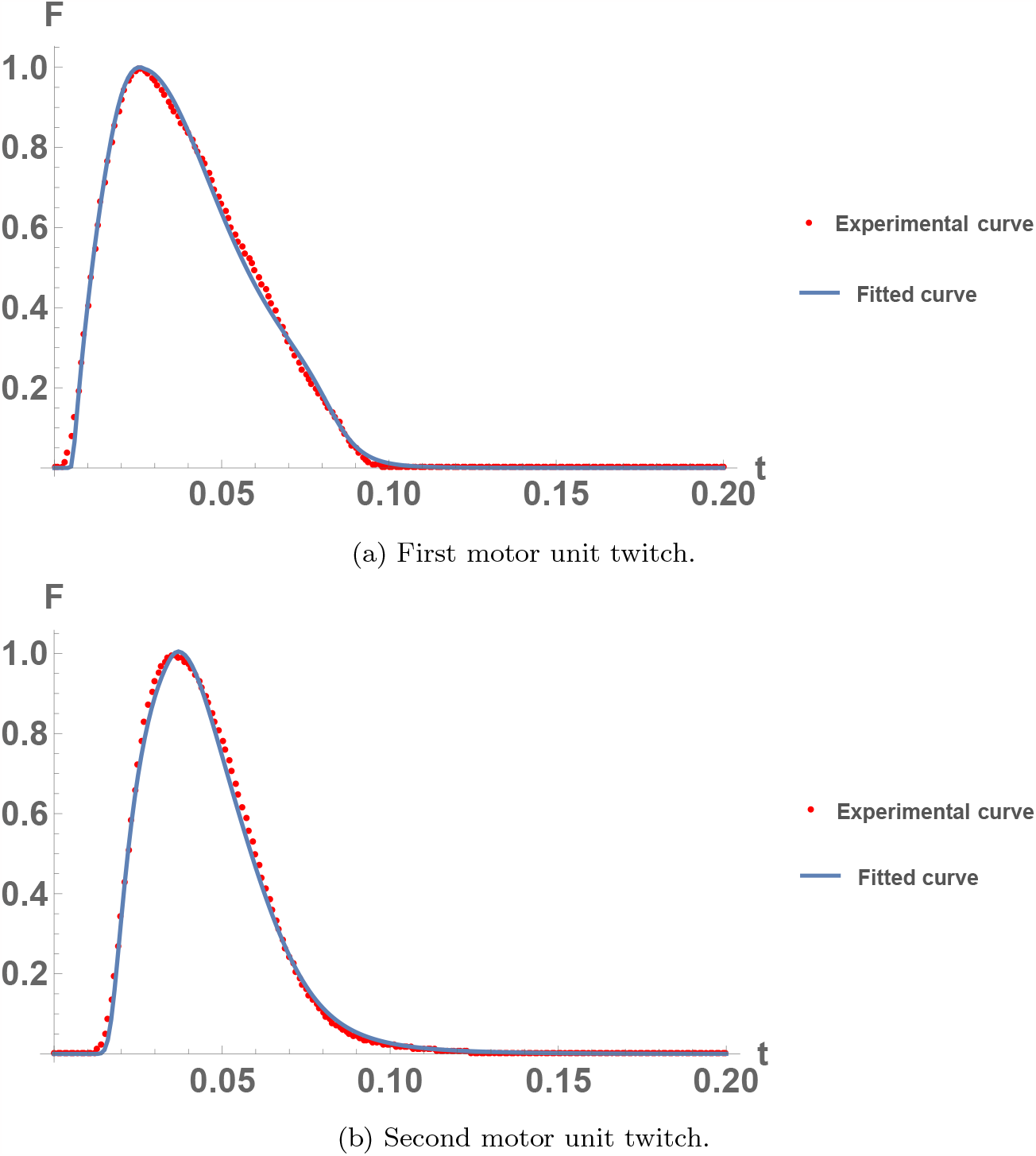
Comparison between the model solution and experimental twitch data for two different medial gastrocnemius muscle MUs of a rat.

**p** = (*k*_*r*_, *k*_*−r*_, *k*_*o*_, *k*_*c*_, *q*_1_, *q*_*−*1_, *q*_2_, *q*_3_, *R*_*T*_, *E*_*T*_, *a*_*quant*_, *k*_*p*_, *C, S, k*_20_, *k*_3_, *k*_4_, *k*_5_, *F*_0_, *µ*_*s*_, *B, A*_1_, *A*_2_) are parameters in the considered model (1), (4), (6) and *F* (*t*_*i*_; **p**) is the solution of the mathematical model (1), (4), (6) for parameters **p** at time *t*_*i*_. We solve the minimization problem

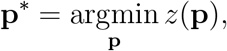

using the Nelder–Mead method [43, 44], built in Wolfram Mathematica. The computation of *F* (*t*; **p**) for any given values of **p** is done by using a fourth-order Runge–Kutta method [45, 46] for the numerical solution of the corresponding ODE systems (1), (4), and (6).

For the first MU twitch, we obtain the result, shown in Fig. 4a, for the parameters, given in Table 1. For the second MU twitch, we obtain the result in Fig. 4b, for the parameters, given in Table 2. As one can see, there is an excellent agreement between the experimental data and the solution of the proposed mathematical model.

**Table 1:**
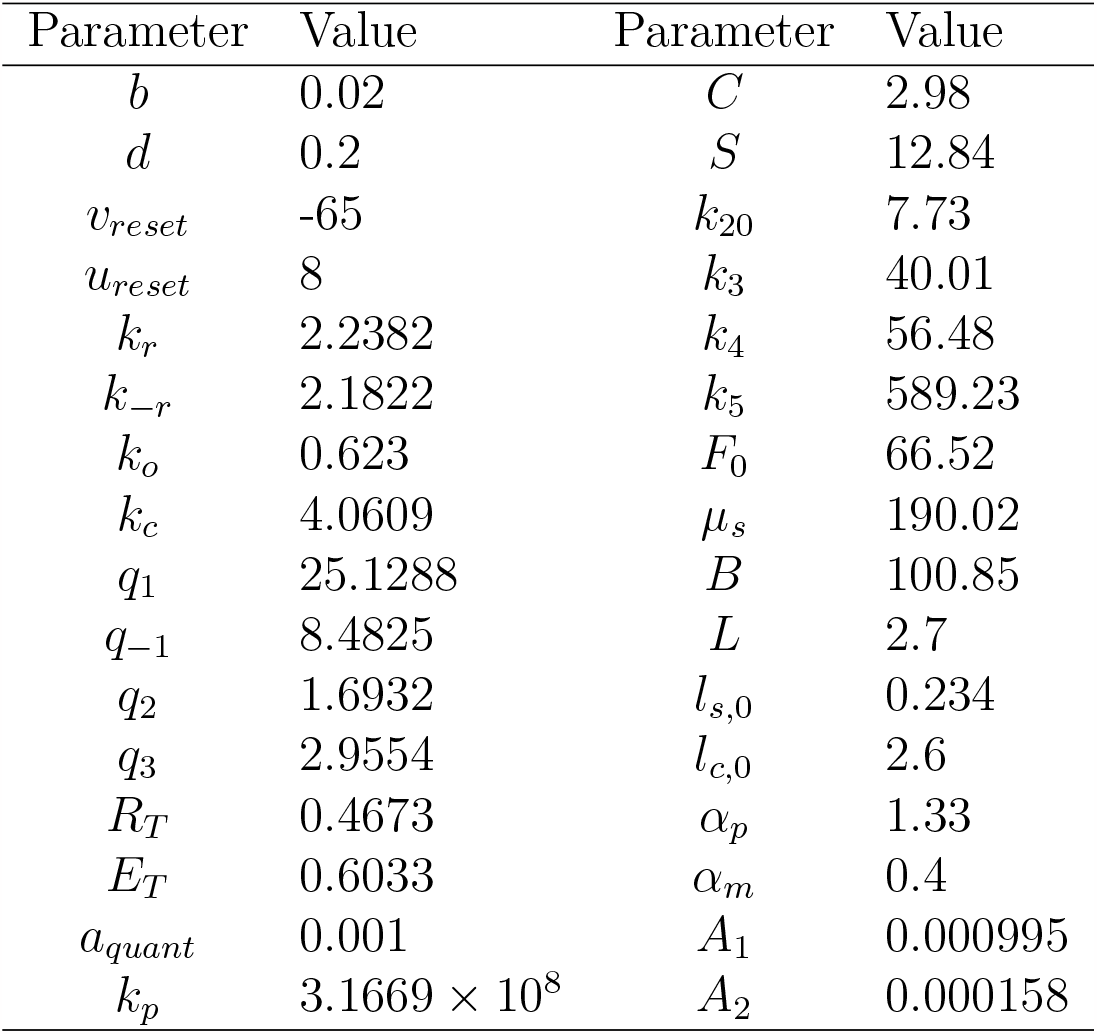
Obtained model parameters for the first MU twitch, see Fig. 4a.

| Parameter | Value | Parameter | Value |
| --- | --- | --- | --- |
| $b$ | 0.02 | $C$ | 2.98 |
| $d$ | 0.2 | $S$ | 12.84 |
| $v_{reset}$ | -65 | $k_{20}$ | 7.73 |
| $u_{reset}$ | 8 | $k_3$ | 40.01 |
| $k_r$ | 2.2382 | $k_4$ | 56.48 |
| $k_{-r}$ | 2.1822 | $k_5$ | 589.23 |
| $k_o$ | 0.623 | $F_0$ | 66.52 |
| $k_c$ | 4.0609 | $\mu_s$ | 190.02 |
| $q_1$ | 25.1288 | $B$ | 100.85 |
| $q_{-1}$ | 8.4825 | $L$ | 2.7 |
| $q_2$ | 1.6932 | $l_{s,0}$ | 0.234 |
| $q_3$ | 2.9554 | $l_{c,0}$ | 2.6 |
| $R_T$ | 0.4673 | $\alpha_p$ | 1.33 |
| $E_T$ | 0.6033 | $\alpha_m$ | 0.4 |
| $a_{quant}$ | 0.001 | $A_1$ | 0.000995 |
| $k_p$ | $3.1669 \times 10^8$ | $A_2$ | 0.000158 |

Let us note that the experimental data were normalized and one needs to rescale back by the maximum force the values of *µ*_*s*_, *F*_0_ and *F* in order to obtain the original experimental data.

For the set of parameters in Table 1, the solutions for *a, r*_*o*_, *k*_1_, *f*_*b*_ and *c*, shown in Fig. 5, were obtained. The numerical results succeed to restore the sequence of events that result in MU force generation—when ACh is released in the NMJ the concentration of open channels, *r*_*o*_, increases. Once the stimulus is on, i.e., *k*_1_ *>* 0 holds true, the concentrations *c* and *f*_*b*_ gradually rise, which leads to the obtained MU twitch. An analogous result can be obtained for the parameters in Table 2.

**Figure 5:**
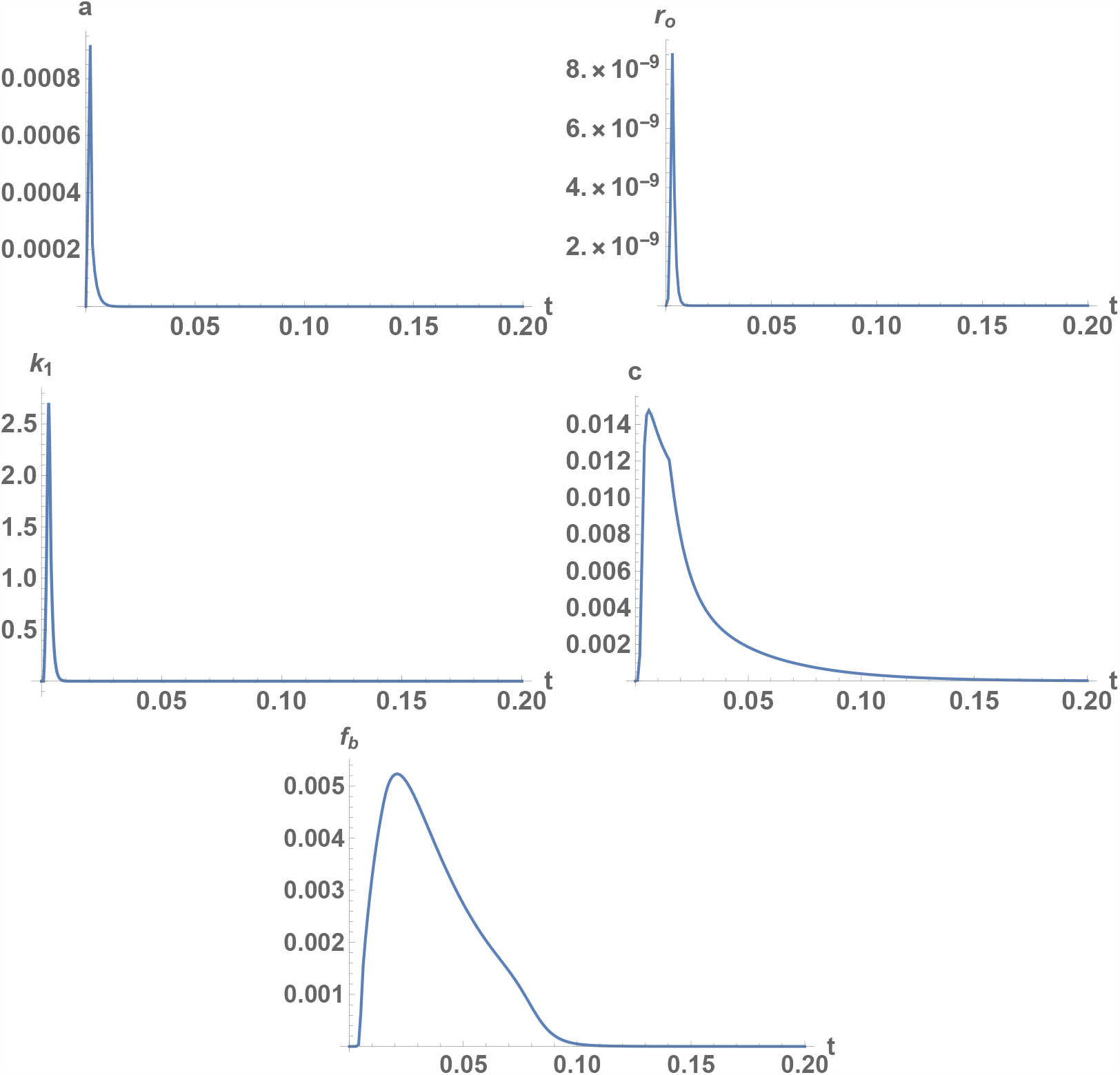
Model solutions, corresponding to the parameters in Table 1, for ACh concentration in the NMJ, *a*; open ACh receptors on the end-plate, *r*_*o*_; rate of calcium release from the SR, *k*_1_; free calcium ions, *c*; bound CF sites, *f*_*b*_.

**Table 2:**
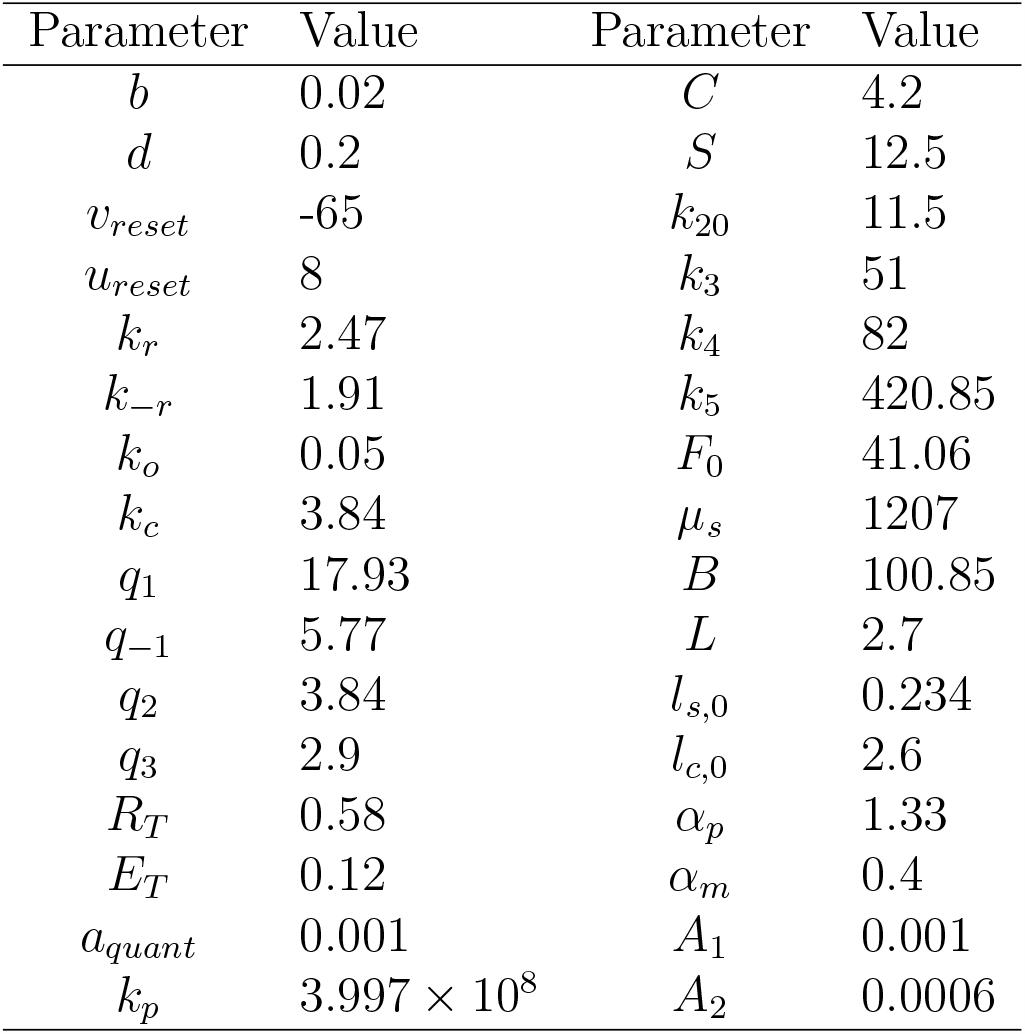
Obtained model parameters for the second MU twitch, see Fig. 4b.

## 4. Sensitivity analysis. Effect of varying the model parameters, related to the NMJ properties, on the characteristics of the twitch

### 4.1. Characteristic parameters of a single twitch

The MU twitch has a characteristic shape, which has been described analytically with various degrees of detailedness. In [1], Raikova et al. have shown that the shape of the twitch can be well described by the following six parameters, see Fig. 6:

**Figure 6:**
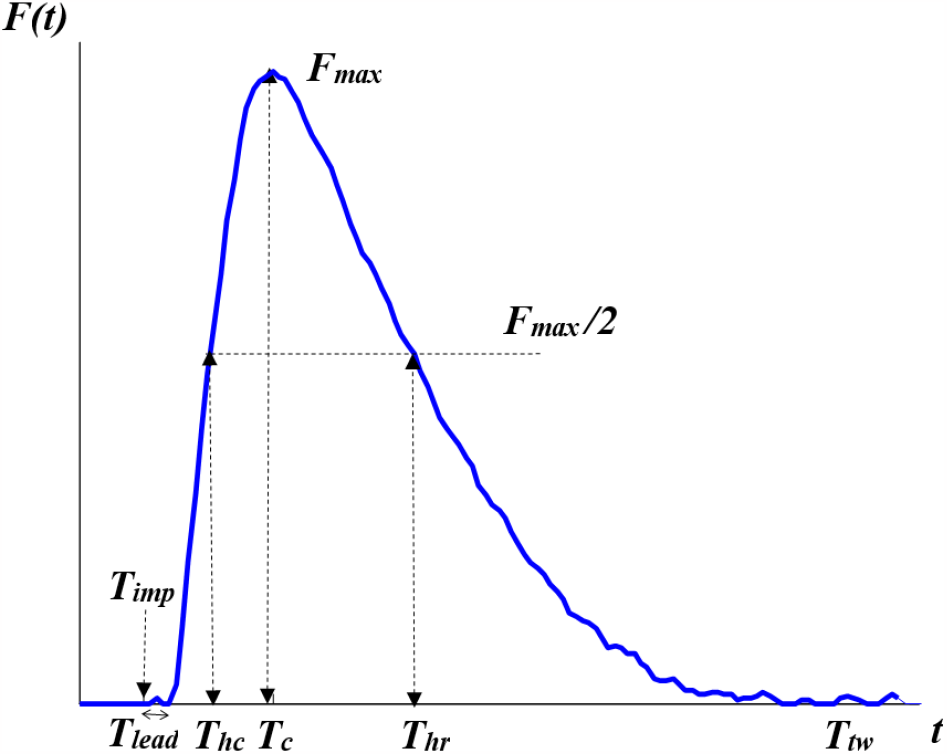
Characteristic parameters for the shape of a twitch—maximal force, *F*_*max*_; time of applied stimulus, *T*_*imp*_; lead time, *T*_*lead*_; contraction time, *T*_*c*_; half-contraction time, *T*_*hc*_; half-relaxation time, *T*_*hr*_; time of the total twitch, *T*_*tw*_. The blue line is constructed from a real experimentally measured twitch of rat medial gastrocnemius muscle MU.

*F*_*max*_—the maximal force;

*T*_*lead*_—the lead time, i.e., the time between the action potential (denoted below by *T*_*imp*_) and the start of force development;

*T*_*c*_—the contraction time, i.e., the time between the start of force development and the moment when the force reaches its maximum;

*T*_*hc*_—the half-contraction time, i.e., the time between the start of the contraction and the moment when the force reaches half of its maximum;

*T*_*hr*_—the half-relaxation time, i.e., the time between the start of the mechanical response and the moment during relaxation period when the force reaches half of its maximum;

*T*_*tw*_—the time between the start and the end of the contraction (defined as the time when MU force decreases to *F*_*max*_*/*1000).

Then, the shape of the twitch, i.e., the force *F* (*t*), can be expressed in terms of the following formula:

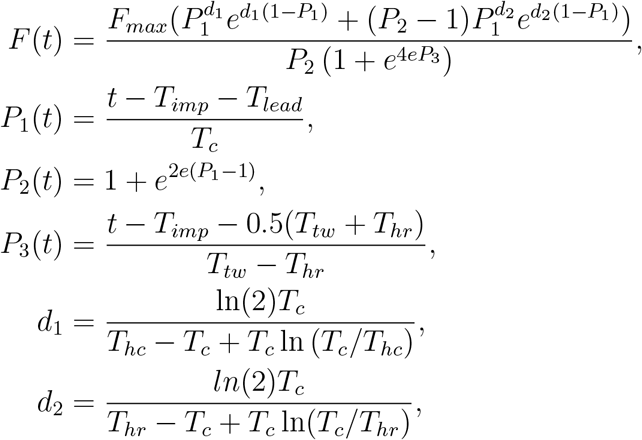

for *T*_*imp*_ + *T*_*lead*_ *< t < T*_*tw*_, *T*_*hc*_ *< T*_*c*_ *< T*_*hr*_ *< T*_*tw*_, where *t* is time. If *t < T*_*imp*_ + *T*_*lead*_ or *t > T*_*tw*_, then one defines *F* (*t*) = 0.

We are now interested in studying how varying the different parameters in our model (1), (4), (6), related to the NMJ properties, affects the characteristics of the twitch shape, in particular, *F*_*max*_, *T*_*c*_, *T*_*hc*_, *T*_*hr*_, *T*_*tw*_. Let us note that the numerical results in this section are obtained by using the parameters in Table 1, varying only a single parameter for each experiment.

### 4.2. Varying the end-plate properties

#### 4.2.1. Effect on the twitch shape

Let us first focus on the effect of the end-plate properties. These properties can be changed due to illnesses, such as, e.g., Myasthenia gravis, which is an autoimmune NMD caused by autoantibodies against the acetylcholine receptor [9, 47].

That is, we are interested in how varying each of *k*_*r*_, *k*_*−r*_, *k*_*o*_, *k*_*c*_ in our model affects the shape of the MU twitch. The results are given in Fig. 7. In Fig. 7a, we show the effect on the maximum force, *F*_*max*_, and in Fig. 7b, we present the effect on the contraction time, *T*_*c*_, half-contraction time, *T*_*hc*_, half-relaxation time, *T*_*hr*_, and the total duration of the twitch, *T*_*tw*_.

**Figure 7:**
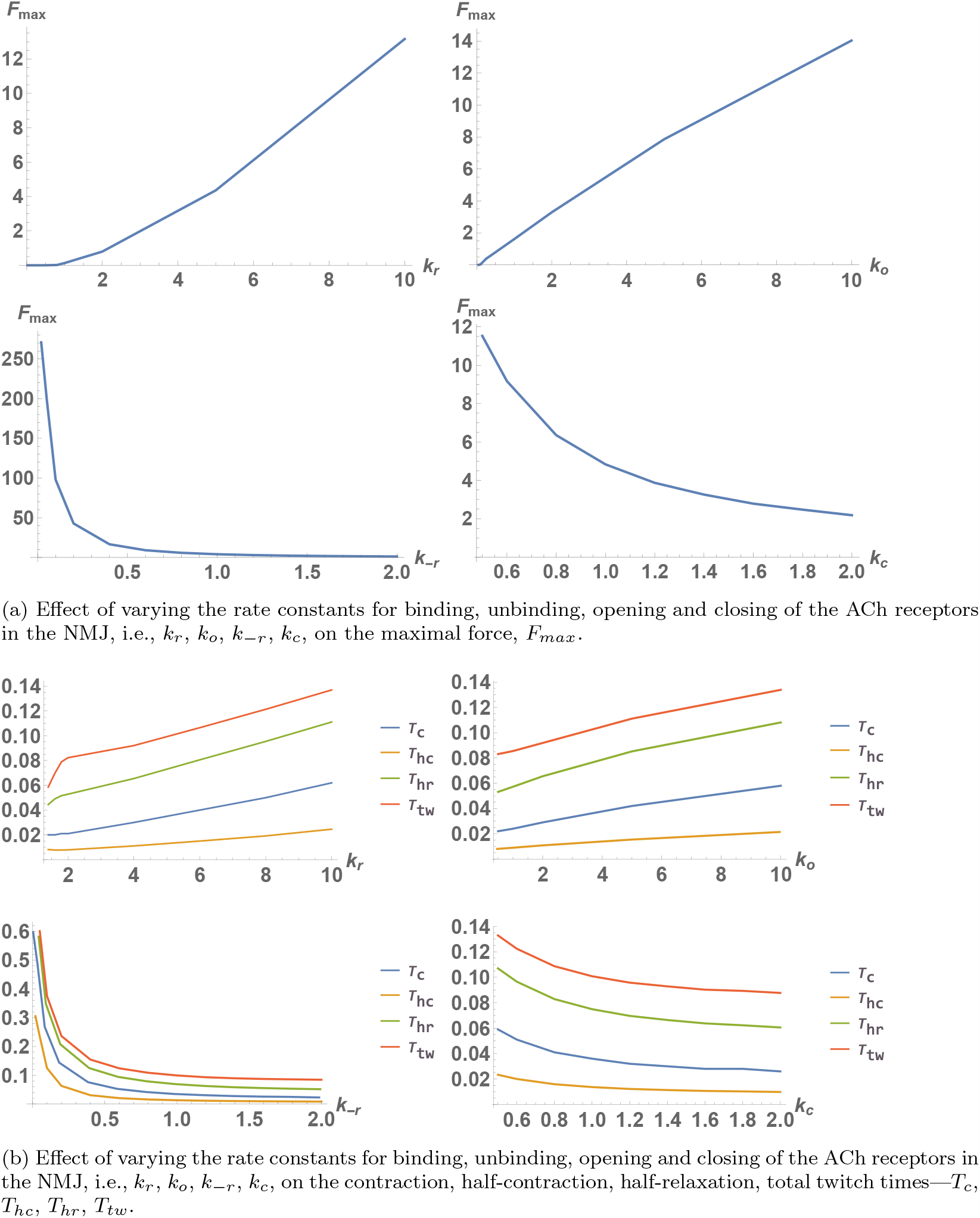
Effect of varying the rate constants, corresponding to the ACh receptors in the NMJ, on the characteristic parameters of the MU twitch.

As can be seen from Fig. 7a, an increase in *k*_*r*_ or *k*_*o*_ leads to an increase in the maximum force. The increase in *k*_*r*_ means that the ACh binds faster to the receptors, which leads to a higher concentration *r*_*o*_ and stronger stimulation of the muscle cell. The higher values of *k*_*o*_ lead to quicker opening of double-bound receptors and again stronger stimulation. As can be expected, an increase in *k*_*−r*_ or *k*_*c*_ has the opposite effect.

Let us note that for small values of *k*_*r*_ or *k*_*o*_, the maximum force, *F*_*max*_, is very close to 0. The reason lies in the nature of the activation function *𝒜*(*f*_*b*_), see Fig. 8. Since *f*_*b*_ needs to reach a certain threshold, before a force is generated, this corresponds to a certain threshold in *k*_*r*_ or *k*_*o*_, as well. For example, as can be seen from Fig. 8, *k*_*r*_ needs to become about 1 so that the maximum of *f*_*b*_(*t*) reaches the threshold of the activation function. Indeed, in Fig. 7a we can see that *F*_*max*_ is indistinguishable from 0 for *k*_*r*_ *<* 1. Similar analysis can be done for *k*_*o*_.

**Figure 8:**
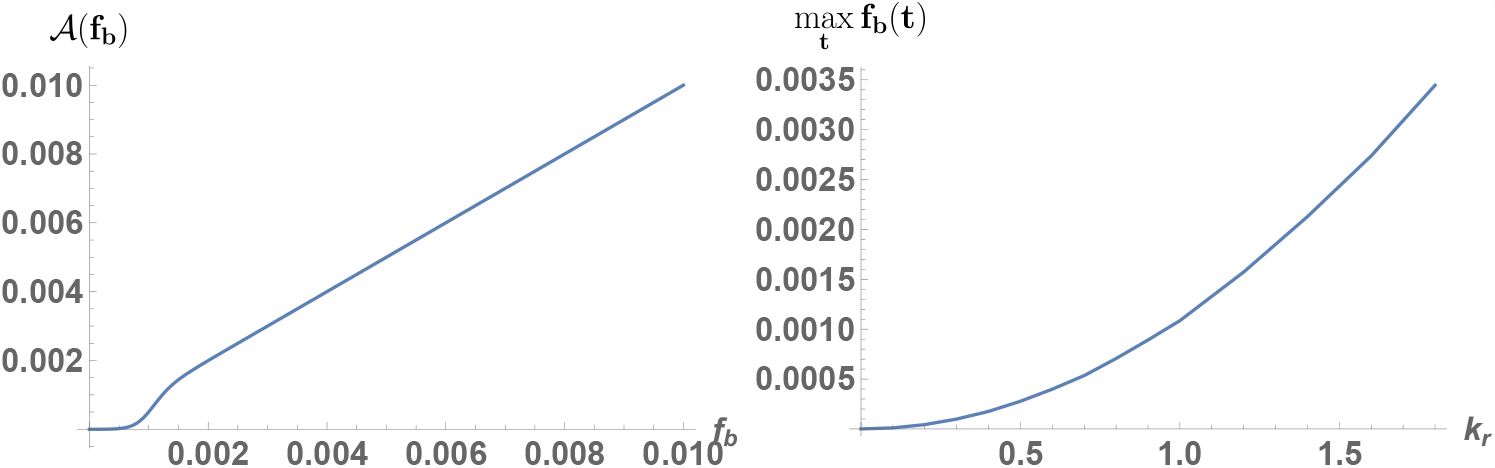
Graph of the activation function *𝒜*(*f*_*b*_) (left) and the maximum of *f*_*b*_ (right) as *k*_*r*_ is varied. The concentration *f*_*b*_ needs to reach a certain threshold before *𝒜*(*f*_*b*_) becomes non-negligible. Therefore, if *k*_*r*_ is very small, i.e., 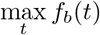 is less than that threshold, the model would predict no generated force.

Besides the maximum of the twitch force, varying the parameters affects significantly its shape as can be seen from Fig. 7b. Firstly, an increase in *k*_*r*_ and *k*_*o*_ leads to an increase in the four characteristic times of the twitch. Furthermore, higher values correspond to an increase of the contraction time *T*_*c*_ relative to the relaxation time (i.e., *T*_*tw*_ *− T*_*c*_). This effect is also illustrated in Fig. 9. Small values of the parameters correspond to quick and weak contraction with relatively long relaxation, while higher values correspond to slow and strong contraction with relatively short relaxation. It seems that the relaxation phase (characterized by *T*_*tw*_ *− T*_*hr*_ and *T*_*tw*_ *− T*_*c*_) is much less affected by those parameters. Again, the effect of *k*_*−r*_ and *k*_*c*_ is the opposite. Let us discuss the effect of those parameters in more detail, because in some sense they have the most direct effect on the MU stimulation and, thus, on the generated force during a twitch.

**Figure 9:**
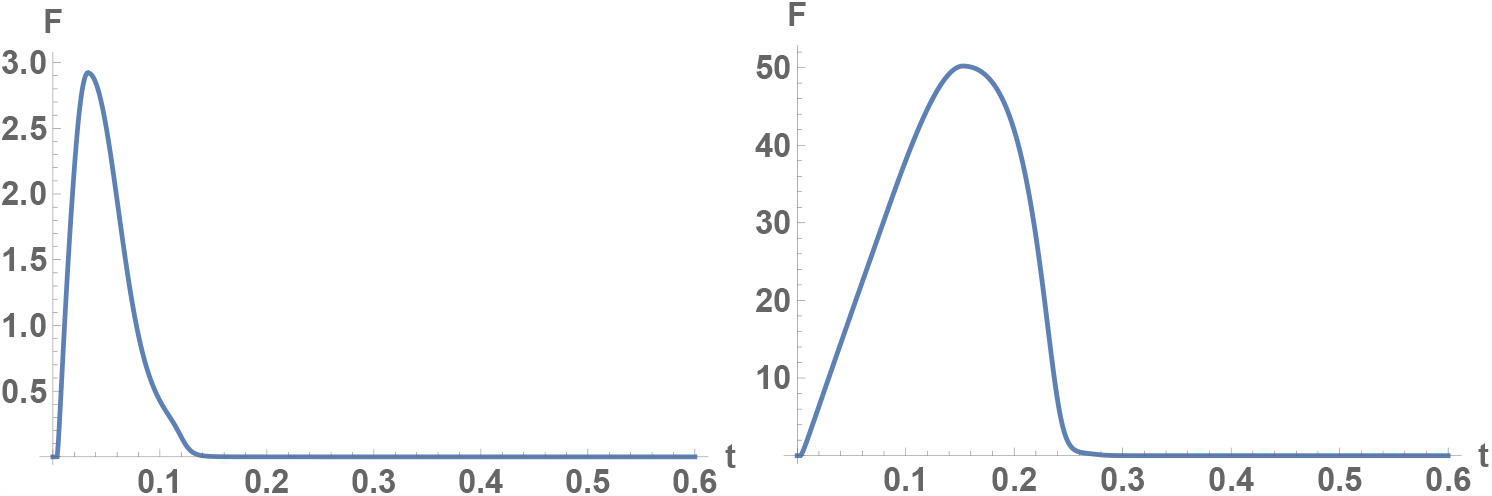
Change of the shape of the twitch as one varies the end-plate properties, calculated for parameters in Table 1, with *k*_*r*_ = 4 (left) and *k*_*r*_ = 30 (right). Increase in *k*_*r*_ leads to longer and stronger contractions and the contraction phase becomes longer than the relaxation phase.

Generally speaking, the variation and the interplay between the rate constants, corresponding to the end-plate of the muscle cell, can have a visible impact on the shape of the twitch. Larger values of *k*_*r*_, *k*_*o*_ or, correspondingly, smaller values of *k*_*−r*_, *k*_*c*_ mean that the muscle cell is being stimulated for longer times, which means that the contraction time increases as does the corresponding maximal force. Since this is related to the longer time of the muscle cell being stimulated, this changes the shape of the twitch from having a steep contraction with slower relaxation (see Fig. 9 on the left) to a situation of a slow (but strong) contraction with relatively quick relaxation (see Fig. 9 on the right).

It is further interesting to make distinction between the effect of the different parameters on the MU twitch shape by comparing twitches, having the same maximum, obtained by varying each of the parameters *k*_*r*_, *k*_*o*_, *k*_*−r*_, *k*_*c*_. Let us set *F*_*max*_ to be 10 and vary successively each of the parameters until we obtain a twitch, having such a value of *F*_*max*_. The results are depicted in Fig. 10. From those graphs it is further evident that varying the endplate properties has a much more distinctive effect on the contraction phase. The contraction phase has a different slope, if we want to obtain the same maximum, by varying the different parameters. The quickest increase, i.e., the most direct effect, is observed by increasing *k*_*o*_. Varying the rest of the properties can also lead to such magnitude of the maximum force, but the contraction times increase. On the other hand, the relaxation phases of all four twitches seem to be very similar.

**Figure 10:**
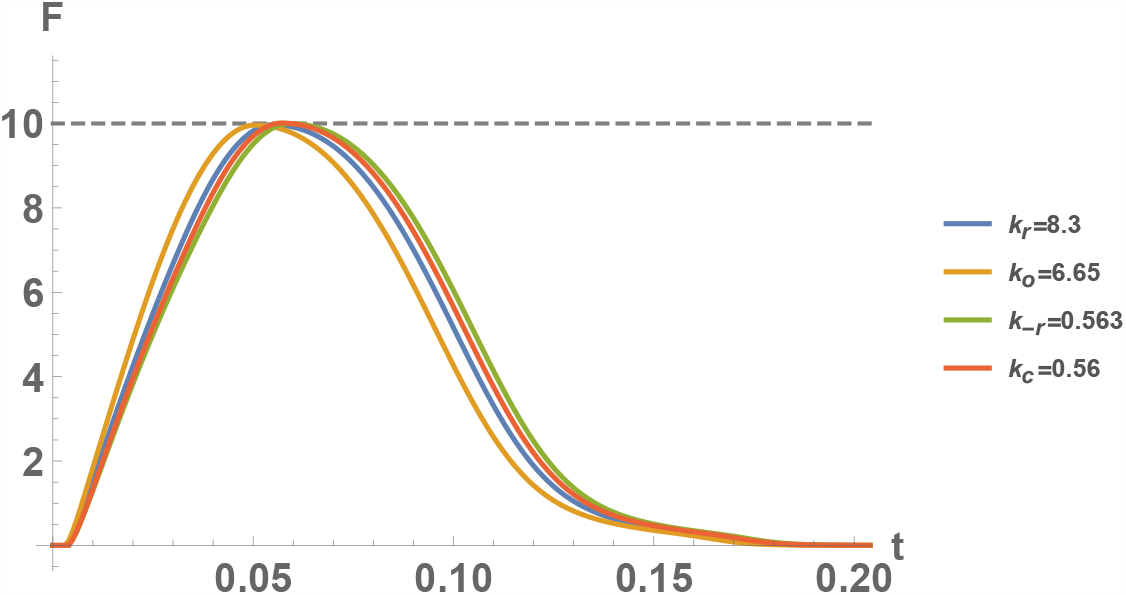
Model solutions *F* (*t*) for parameters, given in Table 1, with one rate constant, corresponding to the ACh receptors state, changed. The respective value of *k*_*r*_, *k*_*o*_, *k*_*−r*_ or *k*_*c*_ is chosen, such that *F*_*max*_ = 10. Each parameter has a distinctive effect on the shape of the twitch.

### 4.2.2. Asymptotic behaviour of the model solutions

For the sake of mathematical completeness, we are also interested in the asymptotic behaviour of the model solutions as *k*_*o*_ and *k*_*r*_ get very large or as *k*_*c*_ and *k*_*−r*_ get very close to 0. Even though such large values, as the ones considered in this subsection, probably have no biological significance for a number of reasons, the results might turn out to be useful to better understand the behaviour of the model.

Useful information to that end can be obtained by considering only the reaction scheme (2), i.e., looking for quasi-stationary states, assuming that the reactions (3) happen on a slower time-scale, as will be further illustrated on the basis of numerical experiments. Then, let us consider the ODE system

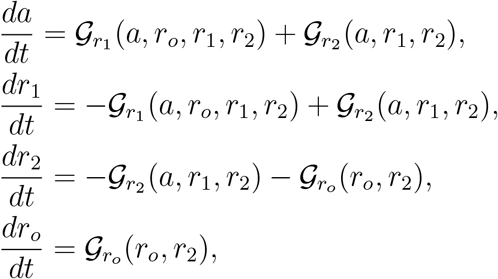

where we have used the notation, introduced in (5). The latter system has a one-parameter family of equilibria, having the form

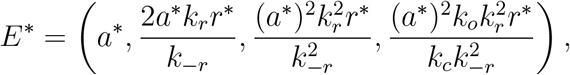

where *a*^*∗*^ is an arbitrary positive number and

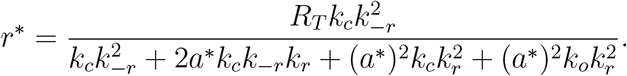

Let us note that the equilibria depend on the rate constants in the form of their ratios *k*_*o*_*/k*_*c*_ and *k*_*r*_*/k*_*−r*_. Furthermore:

if *k*_o_/*k*_c_ → ∞, then *E*∗(a∗,0,0,*R*_*T*_);

if *k*_r_/*k*__r_ → ∞,then 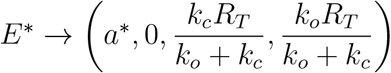

if *k*_o_/*k*_c_ → ∞,then 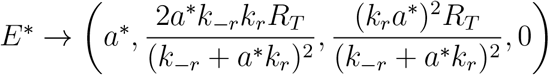

if *k*_r_/*k*__r_ → ∞ (*a*,0,0,0).

Obviously the equilibria in the first two cases might only exist if there is sufficient ACh to open (or at least double-bind to) all receptors with *a*^*∗*^ *>* 0 remaining in free form. If this is not the case, the system will only “strive” for this equilibrium until the ACh is depleted and with binding processes being much faster, the system will reach a quasi-stationary state. This situation can be formalized by considering the approximating reaction schemes:

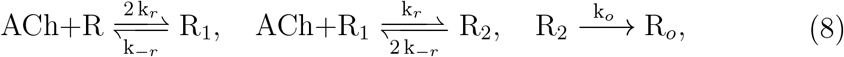

when *k*_0_ is much larger than *k*_*c*_, and

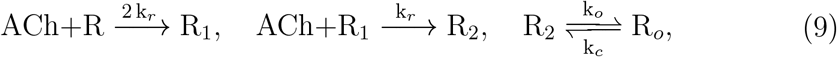

when *k*_*r*_ is much larger than *k*_*−r*_. It is easy to show that (8) has a one-parameter family of zero free ACh equilibria of the form 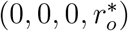 and the reaction scheme (9) has a two-parameter family of zero free ACh equilibria of the form 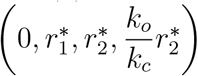 . In both cases, after ACh gets depleted, the system reaches a stationary state with a positive concentration of the open ACh receptors (which is a quasi-stationary state for the original system in the limiting cases). Having this in mind, we are ready to discuss the asymptotic behaviour of the model solutions as *k*_*o*_*/k*_*c*_ *→ ∞* and *k*_*r*_*/k*_*−r*_ *→ ∞*.

Let us first discuss the results for *k*_*o*_*/k*_*c*_ *→ ∞*. We shall, e.g., take large values of *k*_*o*_ with *k*_*c*_ being fixed and numerically solve the model (1), (4), (6) to illustrate such a situation. If we choose some very large value for *k*_*o*_, the dynamics of the ACh receptors is depicted in Fig. 11a. After a quick increase in the ACh concentration and the values of *r*_1_, the solutions quickly tend to a state, where only *r*_*o*_ has significant values with the rest, being close to 0.

**Figure 11:**
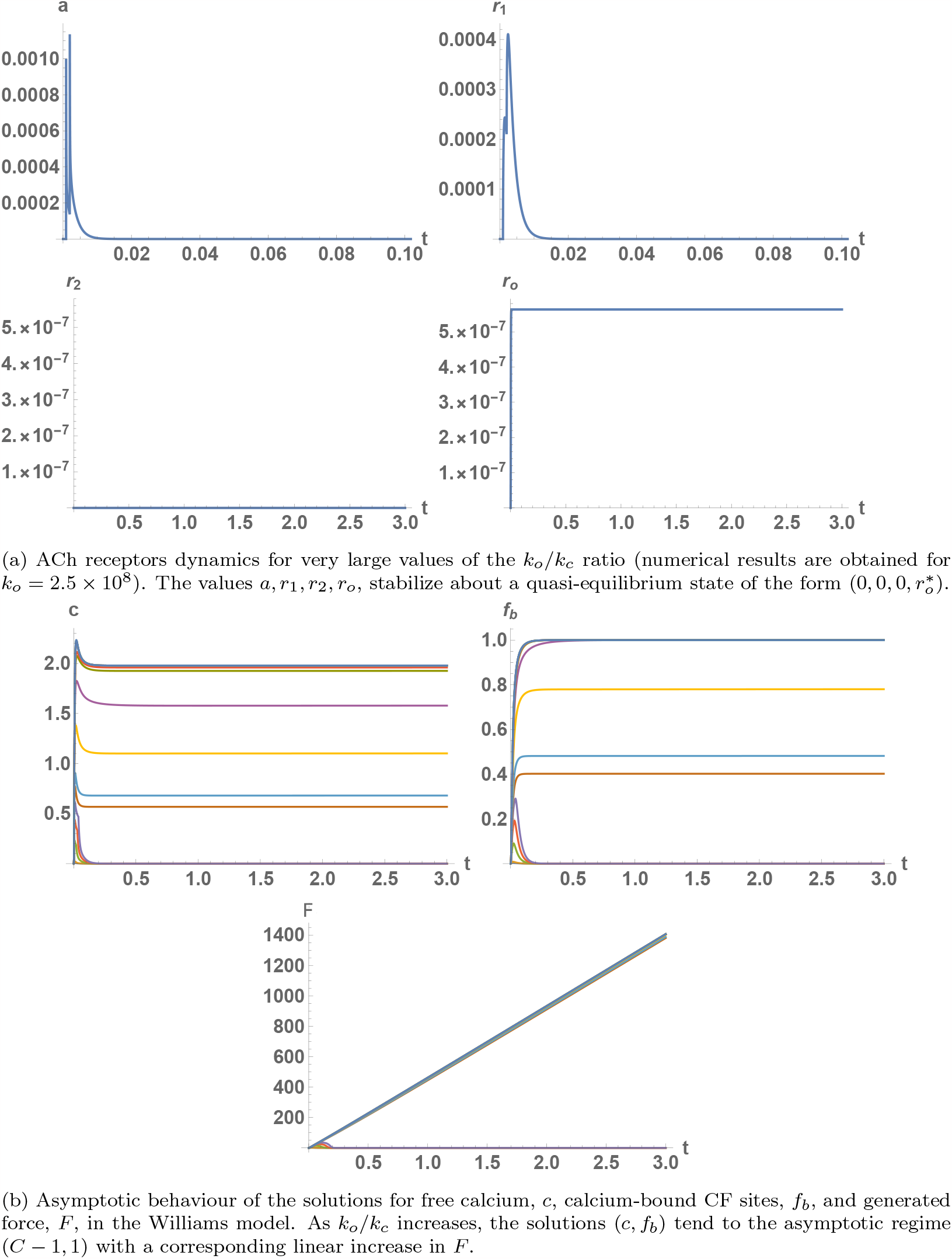
Asymptotic study of the behaviour of the model solutions for large values of the ratio *k*_*o*_*/k*_*c*_.

As *k*_*o*_ increases gradually to such large values, we obtain the behaviours, presented in Fig. 11b. With the increase of *k*_*o*_, there appears a quasistationary state for *c* and, thus, in *f*_*b*_, which leads to a close to linear increase in the resultant force for the observed time period. With the ACh receptors being open, the calcium dynamics inside the muscle cell for large values of *k*_*o*_ follows the quasi-stationary regime of *k*_1_ *≈ const, k*_2_ = 0, studied by Ivanova and Ivanov in [41]. In this case, *c* and *f*_*b*_ tend to *C −* 1 and 1, respectively.

In Fig. 12, we present the results for large values of the ratio *k*_*r*_*/k*_*−r*_. The asymptotic behaviour for *F* (*t*) is similar, but there are substantial differences how this asymptotic behaviour is obtained. As one can see in Fig. 12a, for very large values of the ratio *k*_*r*_*/k*_*−r*_, after a quick increase in *a*, the values *a, r*_1_, *r*_2_, *r*_*o*_, indeed, stabilize about a quasi-equilibrium state of the form 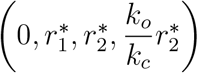 . In Fig. 12b, it is shown that the values *c* = *C −* 1 and *f*_*b*_ = 1 get reached for smaller values of the ratio and those concentrations are retained for gradually increasing periods of time. When we vary *k*_*o*_*/k*_*c*_, the maximum values of *c* and *f*_*b*_ increase much more slowly, but the duration of the stimulus is larger. Therefore, those ratios affect the shape of the twitch in distinctive ways. Increasing *k*_*r*_ leads to a higher maximum of the force, but shorter twitches and vice versa. One can also make such an observation from Fig. 7a. The profile of the first graph is convex, while the second graph is concave.

**Figure 12:**
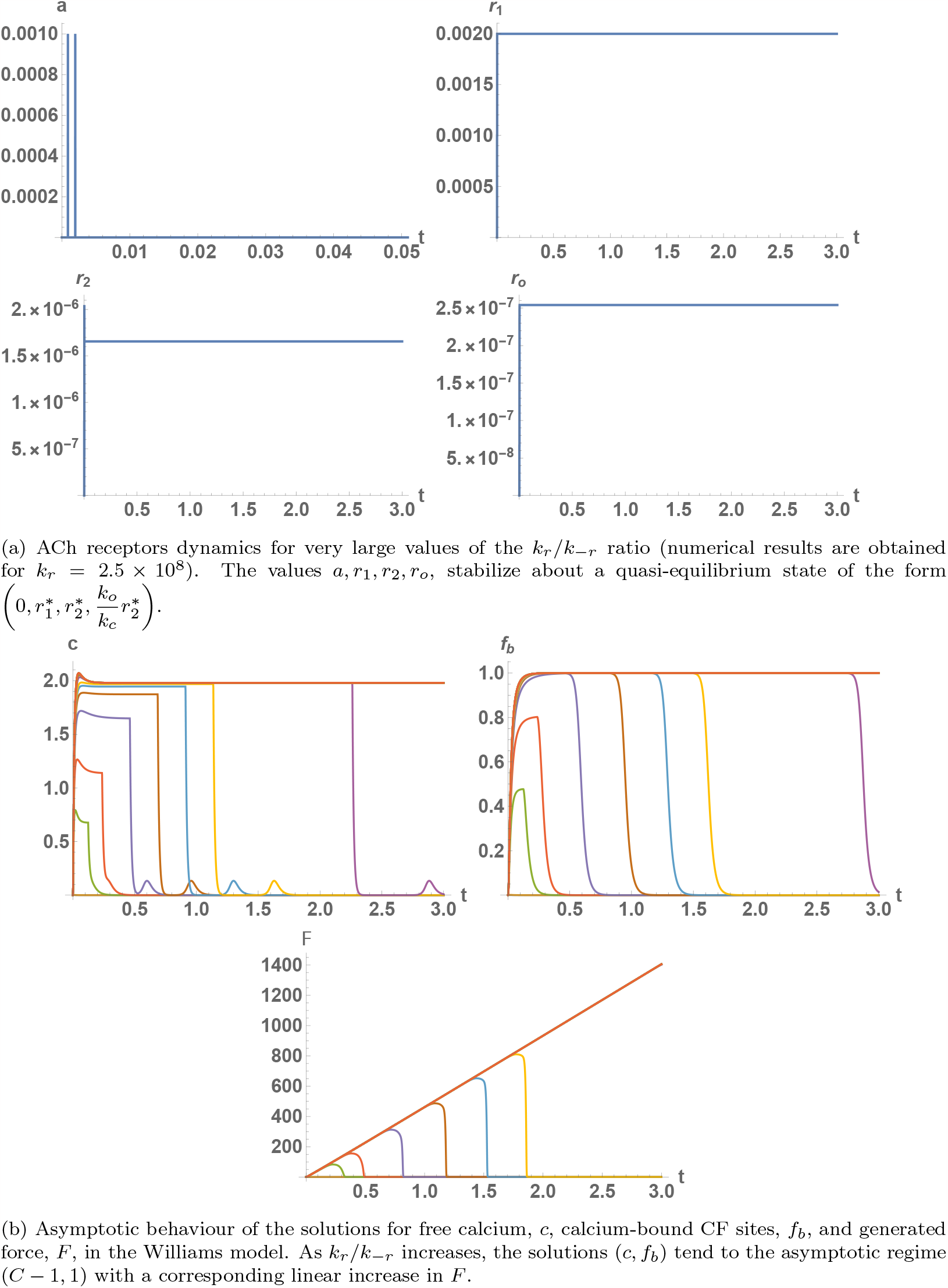
Asymptotic study of the behaviour of the model solutions for large values of the ratio *k*_*r*_*/k*_*−r*_.

### 4.3. Varying the ACh rate constants in the NMJ

Let us now consider the effect of chemical reactions in the NMJ, see the reaction scheme (3), on the shape of the MU twitch, i.e., we shall discuss how varying the parameters *q*_*−*1_, *q*_1_, *q*_2_ changes its properties. Let us note that our experiments have shown that *q*_3_ has very weak effect on the model solutions for the parameter ranges that we consider and, thus, we do not discuss it further in the text. The results for the effect on the maximum force and on the contraction time, half-contraction time, half-relaxation time and the total duration of the twitch are presented in Fig. 13a and 13b. As can be expected, the increase of parameters *q*_1_ and *q*_2_ results in a decrease of *F*_*max*_, since these parameters are related to the catalyzation and breakdown of ACh in the NMJ. The latter leads to lower concentration of ACh and lowers the quanta of ACh, binding to the end-plate receptors.

**Figure 13:**
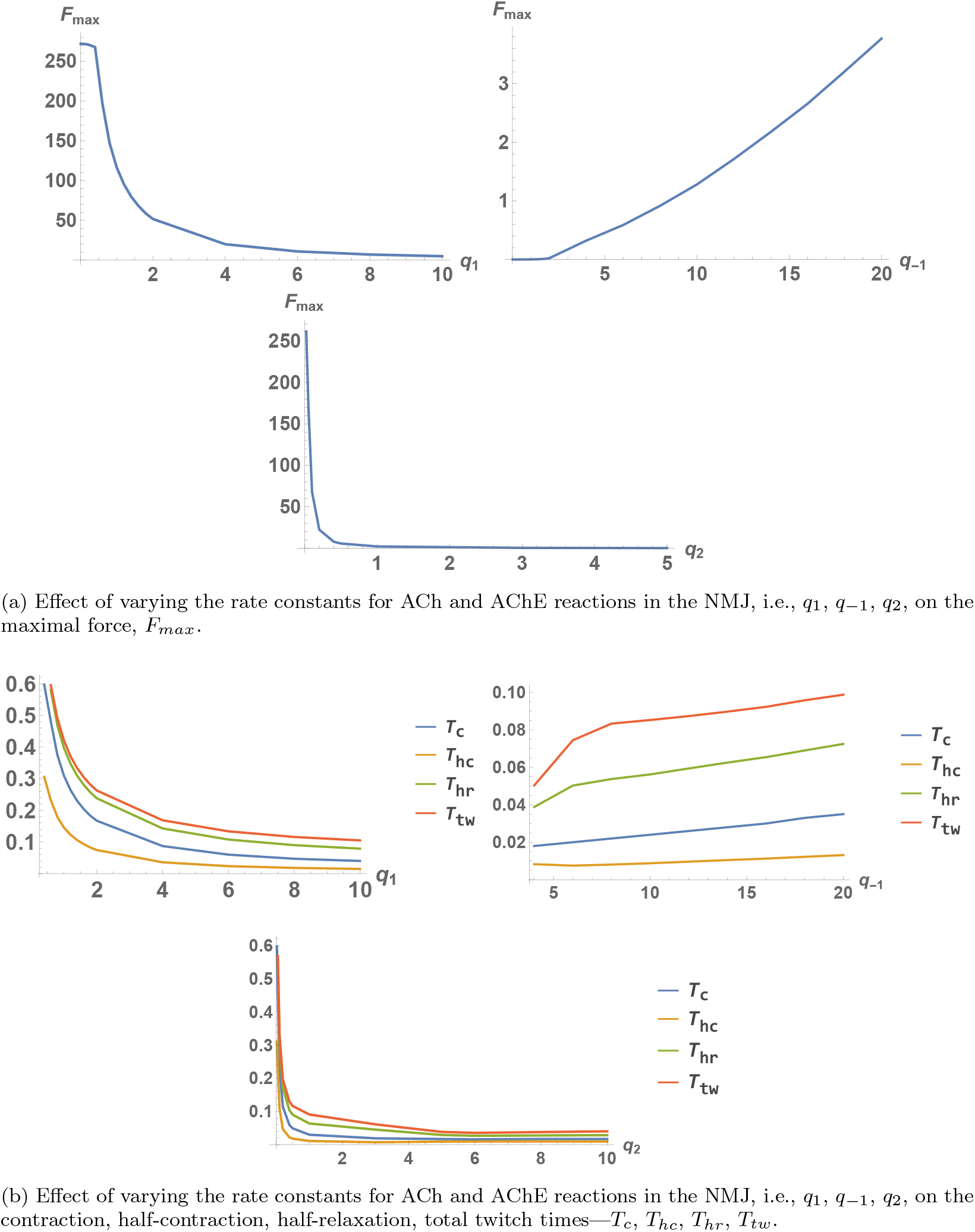
Effect of varying the rate constants, corresponding to the chemical reactions of ACh and AChE in the NMJ, on the MU twitch.

The most significant parameter from this group seems to be *q*_1_. If this parameter is decreased (e.g., using AChE inhibitors [48]), this could significantly increase the force, generated by the MU as well as the duration of the twitch, due to the higher concentrations of ACh, retained in the NMJ. The numerical results suggest that a certain threshold in *q*_1_ exists under which any further decrease in its value would have little effect on the MU force. On the other hand, the reverse reaction, having rate constant *q*_*−*1_ has much lesser effect on the MU twitch for the parameter ranges we consider. As can be seen in Table 1, it is generally several times less than the forward reaction. Thus, its rate should be increased significantly, before a notable change in the twitch is observed. The characteristic parameters of the twitch also seem to be much less sensitive with respect to *q*_2_ within the parameter ranges that we consider. This is natural, because *q*_2_ and *q*_3_ correspond to the rates of chemical reactions, which are only indirectly connected with the ACh concentration in the NMJ and, thus, with the stimulus of the muscle cell.

### 4.4. Varying the values of receptor sites and AChE concentrations

We finally discuss the effect of varying the parameters *R*_*T*_ and *E*_*T*_ . The former can, e.g., be controlled with neuromuscular blocking agents and the latter can, e.g., be associated with AChE deficiency. As can be seen in Fig. 14, a substantial decrease in *E*_*T*_ can lead to a notable increase in the duration of the twitch and its amplitude, due to the prolonged action of ACh in the NMJ and the corresponding stimulation of the end-plate. This is an effect which has been observed in practice, e.g., when using specific anaesthetic drugs on patients with such a disorder [49] as well as in the case of type 1 diabetic neuropathy [50].

**Figure 14:**
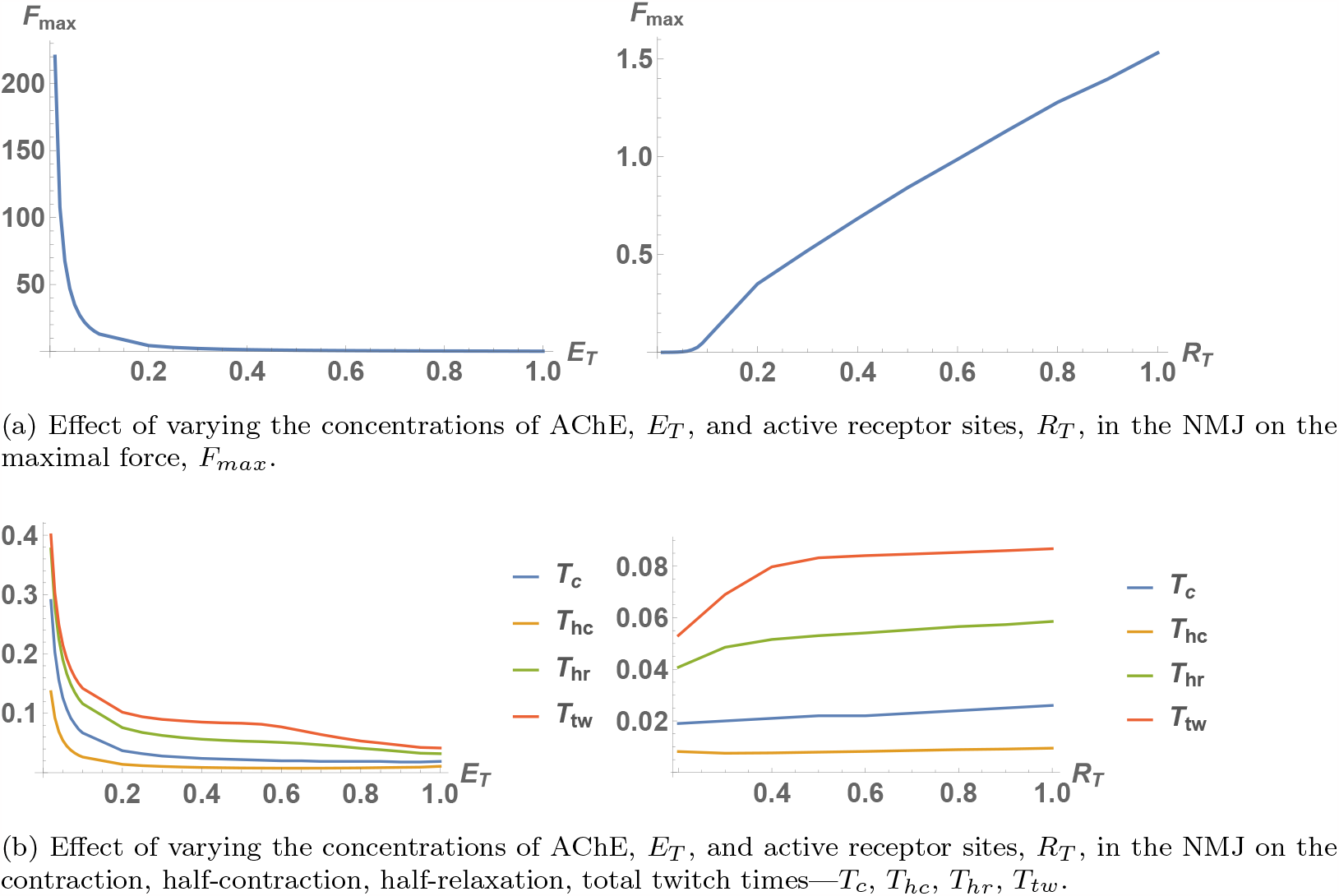
Effect of varying the AChE and active receptor sites concentrations in the NMJ on the MU twitch.

On the other hand, the decrease of active receptor sites *R*_*T*_ (which can be controlled with neuromuscular blocking agents [51]) is associated with a decrease in the amplitude and duration of the twitch. The numerical results show a threshold value of active receptor sites, needed for a twitch to develop.

## 5. Conclusion and discussion

In the present work, we proposed a new integrated mathematical model of neuromuscular activation, coupling known models for nerve impulses, neurotransmitter transport in the NMJ, calcium activity inside the muscle cell and resultant MU force generation. Using this model, we managed to successfully recreate experimental data for twitches of two different MUs, thus, validating its applicability to accurately simulate various MU twitches. Further, we studied how the main parameters, related to the properties of the NMJ, affect the shape of the twitch. It is important to note that the parameters, with respect to which the twitch shape seems to be most sensitive, could be directly related to known NMDs and/or treatments on the neuromuscular system. This is very promising in the sense that in the future the model could successfully be applied to the study of NMDs and could give useful results to that end.

In this direction, as a future work one should make a classification of known NMDs with respect to the model parameters they are related to. Then, a careful analysis for each disease should be done, taking into account the known experimental and clinical data and studying its relation to the results, given by the numerical simulations.

On the other hand, a more extensive collection of MU activities should be simulated, using the model, including various MU twitches as well as tetanic contractions. Thus, one should obtain a collection of parameter sets, corresponding to different possible behaviours of neuromuscular activity.

## List of abbreviations

ACh: acetylcholine
NMD: neuromuscular disease
AChE: acetylcholineesterase
NMJ: neuromuscular junction
CF: contractile filament
ODE: ordinary differential equation
MU: motor unit
SR: sarcoplasmic reticulum

